# Integration of Dominance and Marker *×* Environment Interactions into Maize Genomic Prediction Models

**DOI:** 10.1101/362608

**Authors:** Luis Felipe Ventorim Ferrão, Caillet Dornelles Marinho, Patricio R. Munoz, Marcio F. R. Resende

## Abstract

Hybrid breeding programs are driven by the potential to explore the heterosis phenomenon in traits with non-additive inheritance. Traditionally, progress has been achieved by crossing lines from different heterotic groups and measuring phenotypic performance of hybrids in multiple environment trials. With the reduction in genotyping prices, genomic selection has become a reality for phenotype prediction and a promising tool to predict hybrid performances. However, its prediction ability is directly associated with models that represent the trait and breeding scheme under investigation. Herein, we assess modelling approaches where dominance effects and multi-environment statistical are considered for genomic selection in maize hybrid. To this end, we evaluated the predictive ability of grain yield and grain moisture collected over three production cycles in different locations. Hybrid genotypes were inferred *in silico* based on their parental inbred lines using single-nucleotide polymorphism markers obtained via a 500k SNP chip. We considered the importance to decomposes additive and dominance marker effects into components that are constant across environments and deviations that are group-specific. Prediction within and across environments were tested. The incorporation of dominance effect increased the predictive ability for grain production by up to 30% in some scenarios. Contrastingly, additive models yielded better results for grain moisture. For multi-environment modelling, the inclusion of interaction effects increased the predictive ability overall. More generally, we demonstrate that including dominance and genotype by environment interactions resulted in gains in accuracy and hence could be considered for genomic selection implementation in maize breeding programs.

## 1 Introduction

Nearly a century ago, G. H. Schull proposed the term heterosis to describe the higher performance of “cross-bred” individuals when compared with corresponding inbred or “pure-bred” genotypes [Shull, 1948]. Since his pioneer studies, the development of hybrid varieties has been an integral part of many plant breeding programs resulting in significant gains in global grain production. The clearest example of success has been reported in maize (*Zea mays* L.), which hybrid varieties are now widely adopted, replacing open-pollinated populations. Among the advantages, the better yield and greater uniformity are central features that favored its rapid acceptance by companies and producers [Crow, 1998].

Hybrid vigor, in maize, is traditionally obtained by crossing inbred lines from genetically distinct pools, the so-called heterotic groups. Depending on the stage of the breeding program, selected hybrids can be used for commercial deployment, or to continue the breeding cycle and generate new inbred lines using a strategy known as advanced-cycle pedigree breeding [Lu and Bernardo, 2001]. In this process, an important step for single-cross hybrids is the choice of parental lines that have good combining ability and hence can capitalize this hybrid vigor. Traditionally, selection of promising genotypes relies on phenotypic field records and pedigree data, which is a labor-intensive process. As pointed out by Technow et al. [2014], assuming that a breeding program can generate 1000 lines in each heterotic group per year, the number of potential hybrids to be evaluated in the field is 1 million. Given the difficulty to test all combination in field experimental designs, prediction of hybrid performance is one of the current bottlenecks in maize breeding programs. To circumvent this issue, a contemporaneous alternative is to adapt genomic prediction algorithms – originally proposed to predict the genetic merit of an individual [Meuwissen et al., 2001] – to predict hybrid performance. Much of the optimism of this approach is motivated by the opportunity to reduce the cost and labor involved in field trials and increase the genetic gain.

Since genomic selection (GS) was introduced in plant breeding, its ability to predict the genetic merit has been evaluated in several crops for different traits [de los Campos et al., 2012]. Initial developments focused on additive models and largely overlooked dominance and epistatic effects, despite the fact that several lines of evidence suggest that non-additive models drive the genetic basis of heterosis [Birchler et al., 2010]. This source of genetic variation was neglected for different reasons, including the lack of informative pedigrees, computational complexities related to estimation of dominance effects [Vitezica et al., 2013], the thought that most genetic variance is additive [Hill, 2010] or can be captured with additive parameterizations [Huang and Mackay, 2016], and the fact that even when non-additive effects are included in the models they are not easily partitioned from additive effects [Muñoz et al., 2014]. Nonetheless, recent inclusion of non-additive effects have demonstrated increased prediction accuracies in some traits in animal and plant breeding [Technow et al., 2014, de Almeida Filho et al., 2016, dos Santos et al., 2016, Resende et al., 2017, Dias et al., 2018]

In addition to the source of genetic variability controlling a trait of interest, a second relevant issue to plant breeders is how to manage the challenges of genotype-by-environment (G*×*E) interaction. G*×*E interactions are expressed as changes in the relative performance of genotypes across environments, which can affect the genotype ranking. From a statistical point of view, G*×*E can be modeled as an interaction effect in a two-way ANOVA model, assuming genotypes and environments as main effects [Meyer, 2009]. More recently, genotypic performance across the environments has been studied as correlated traits in a multivariate linear mixed models framework [Smith et al., 2005, Meyer, 2009]. One of the first ideas to accommodate this in the GS context was described by Burgueño et al. [2012]. These authors proposed an extension of the traditional GBLUP method [VanRaden, 2008], where G×E interactions were modeled considering variance-covariance structures with heterogeneity of genetic variance across individual environments and heterogeneity of genetic correlations between pair of environments.

After the referred study, new methods addressing G*×*E interaction were proposed and investigated in several crops, including maize [Burgueño et al., 2012, Cuevas et al., 2016a, Lopez-Cruz et al., 2015, Dias et al., 2018, Fristche-Neto et al., 2018, e Souza et al., 2017, Ferrão et al., 2017]. One particular model introduced by Lopez-Cruz et al. [2015] and improved by Cuevas et al. [2016a] and Cuevas et al. [2016b] - explicitly models the interactions of each marker with the environment (M*×*E interaction). The main advantage of this method is the possibility to decompose marker effects into components that are common and specific across environments, which are concepts related to adaptability and stability in the breeding context [Eberhart and Russell, 1966]. Moreover, extending the M*×*E interaction also provides an opportunity to investigate marker effect individually, which ultimately may shed light on the underlying genetic architecture of traits [Crossa et al., 2016]. Although theoretical and empirical studies have shown the potential of GS in multi-environment trials, it is still an open question how well can these models predict a completely new and unobserved environment [Ferrão et al., 2018]. In the literature, much attention has been paid on prediction in the so-called CV2 scheme - in reference to Burgueño et al. [2012]. This approach mimics the prediction of incomplete field trials, and it is relevant in advanced stages of the breeding, where “soon-to-be-deployed” hybrids have their performance predicted in multiple environments. However, the prediction accuracies obtained in a CV2 scheme, does not reflect the case where breeders want to predict the phenotype performance of a new genotype under an untested environmental condition.

Given the potential of GS to reshape breeding programs, in this study we report the expected response of GS using additive and dominance models for predictions of two important traits in maize: grain yield and grain moisture. We also emphasize the benefits to accommodate G*×*E interactions in GS model, in order to achieve fast and longstanding genetic gains. More generally, we provide a critical analysis about GS implementation in multi-environment trials. Although our work is motivated by prediction in hybrid maize, many of the ideas and results can be applied broadly.

## 2 Material and Methods

The description of the Material and Methods is organized as follow. In Section 2.1, we describe the development of the population used in the experiment, collection of phenotypic data and correction of phenotypes taking into account experimental covariates prior to its use as input in the genomic models. Steps related to genotyping, single nucleotide polymorphisms (SNPs) filtering, genetic diversity and definition of the hybrid genotypes *in silico* were described in Section 2.2. The incorporation of additive and dominance effects in Marker *×* Environment (M*×*E) genomic prediction models are described in Section 2.3. Finally, in Sections 2.4 and 2.5 we describe the computational implementation of the proposed methods and explain how these models were compared in our experiment.

### 2.1 Plant Material

The phenotypic data consisted of grain yield (kg/ha) and percentage of grain moisture. Phenotypic records were collected in 1831 hybrids during three years (2015, 2016 and 2017). Each cycle included a different set of single-cross maize hybrids evaluated in different locations, as presented in Table 1. Hybrids were originated from single crosses between 207 inbred lines from different heterotic groups. All field trials were established in Brazil by the company Helix Seeds, São Paulo, Brazil. The presented analysis considered the collective result from each year as a different environment. Phenotypes were adjusted using linear mixed models for each evaluation cycle (year) in the SELEGEN software [Resende, 2016]. The experimental design was a randomized complete block design with three replicates. For each evaluation cycle, the phenotypic model consisted of locations and blocks considered as fixed effects; and hybrid genotypes treated as independently and identically distributed random effects. The choice of treating genotypes as random effects was made due to the highly unbalanced nature of the data. The Best Linear Unbiased Prediction (BLUP) for each hybrid genotype was used as its phenotypic value in GS models. To compute the empirical phenotypic correlation across the evaluation cycles, we used the commercial hybrids that were used as checks in all environments and years.

**Table 1:** Number of maize hybrids evaluated across three breeding cycles (2015, 2016 and 2017) in eight locations in Brazil.

| Location | State | 2015 | 2016 | 2017 | Latitude | Longitude | Altitude |
| --- | --- | --- | --- | --- | --- | --- | --- |
| Location 1 (IP) | MG | 347 | - | - | 18°38'30" S | 49°53'06" W | 415 |
| Location 2 (PM) | MG | 470 | 580 | 800 | 18°45'17" S | 46°38'28" W | 889 |
| Location 3 (NM) | MT | 618 | 750 | 925 | 13°48'55" S | 56°05'36" W | 475 |
| Location 4 (SE) | PR | 512 | 585 | 714 | 23°09'74" S | 50°58'74" W | 379 |
| Location 5 (SO) | MT | 469 | 580 | 800 | 12°32'00" S | 55°42'00" W | 365 |
| Location 6 (PA) | PR | - | 158 | 97 | 24°16'38" S | 53°49'388" W | 310 |
| Location 7 (PL) | MT | - | 231 | 123 | 15°30'00" S | 54°35'00" W | 668 |
| Location 8 (CN) | MT | - | - | 79 | 13°39'32" S | 53°53'34" W | 550 |
| **Total** |  | **618** | **750** | **925** |  |  |  |

### 2.2 Genotyping and *in silico* crossing

The inbred lines were genotyped using the maize 500k Affymetrix chip. Raw data was filtered, removing SNPs with the following quality control parameters: (i) all markers with missing values, (ii) all markers with any heterozygous calls in the inbred lines, (iii) markers with a minor allele frequency (MAF) < 0.02. This selection process resulted in a set of 24,758 filtered markers. As a first assessment, prediction accuracies in a single location using full and filtered marker set resulted in very similar results (data not shown). The hybrid genotypes were inferred by combining one allele from each of the respective parental lines. In summary, parental bi-allelic marker loci were encoded as *x*_{*AA,BB*}_ ⊂ {0, 2} in the maize lines. When both parental loci were the same genotype, the corresponding hybrid genotype was encoded as homozygous, with the same genotype as their parents. Heterozygous were encoded as *x*_{*AB*}_ ⊂ {1}, when both parents were homozygous for different alleles.

A genomic relationship matrix (**G**) based on the filtered markers was computed, using the respective equation VanRaden [2008]: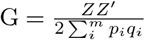 where *p* and *q* are the allele frequencies at locus *i*, and **Z** (originally coded as 0,1,2) is the matrix of centered markers. In order to examine the population structure and diversity within the set of parental lines, we performed a principal component analysis (PCA) applied on the resulting **G** matrix. The first two principal components were considered to represent the population stratification. Individuals were assigned to groups by a k-means clustering approach. Appropriate cluster number was determined by plotting k-values from 1 to 10 against their corresponding within-group sum of squares (SSE).

### 2.3 Statistical Models

Here, we assumed each evaluation cycle (years) – adjusted for the different locations and experimental design – as a different environment. Three statistical approaches to address M*×*E interactions were considered. Firstly, we refer as **single-environment (SE)** the regression of phenotypes on markers separately in each environment. The **across-environment (AE)** method addressed a combined analysis of years, assuming that marker effects are constant across the environments and ignoring the genotype by environment interaction. Finally, the **multi-environment (ME)** method was the M*×*E interaction model that accounted for common and specific marker effects across the environments. All the aforementioned approaches are, in itself, not new and they were previously reported by Lopez-Cruz et al. [2015]; Crossa et al. [2016]; Cuevas et al. [2016a] and Cuevas et al. [2016b]. Compared with these studies, our work makes contribution by expanding the M*×*E model to include dominance effects. This following section presents details about these models.

#### 2.3.1 Single-environment (SE) model

This method fits a Gaussian linear regression, where a pre-adjusted phenotype vector is regressed on a set of markers in each environment. The importance to incorporate marker effects with dominance effects was investigated considering three versions of the SE models, referred as SE additive (Model 1); SE dominance (Model 2); and SE additive+dominance (Model 3). These statistical models are given, respectively, by:

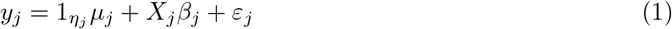

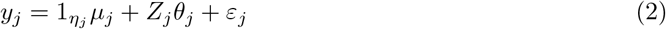

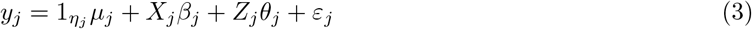

where *y_j_* is the response vector containing *η_j_* pre-adjusted phenotypic values, 1_*ηj*_ is a vector of ones, *µ_j_* is an intercept of the *j* th environment; *X_j_* and *Z_j_* are design matrices relative to additive and dominance effects, respectively. These matrices are representing the allelic state of the hybrids at *p* genetic markers, *X_j_*where denotes the number of reference alleles at a specific locus in the genome (e.g., coded as 0,1,2), while *Z_j_* indicates if this locus is homozygous (coded as 0) or heterozygous (coded as 1). Similar parametrization has been used in plants [Muñoz et al., 2014, Fristche-Neto et al., 2018, Werner et al., 2018] and animal studies [Toro and Varona, 2010]. *β* and *θ* and are *p*-vectors of (unknowns) additive and dominance marker effects, respectively. Following the standard assumptions of the GBLUP model, additive and dominance marker effects were assumed to be independent of each other and both normally distributed: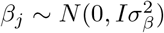 and 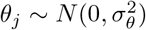. *ε* is a *n*-vector of residual effects with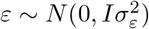.

#### 2.3.2 Across-environment (AE) model

This model assumes that additive and dominance marker effects are the same across environments, that is: *β*_1_ = *β*_2_ = *β*_3_ and *θ*_1_ = *θ*_2_ = *θ*_3_, respectively. The AE additive+dominance model, in a matrix notation, can be written as:

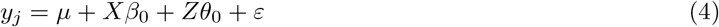

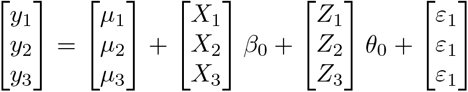

Formally, the main difference compared to the SE approach are matrices and vectors dimensions. Here, all environments are concatenated in order to estimate a common effect. Thus, *y* vector is a *m*-vector of pre-adjusted phenotypes, where *m* is the total number of individuals across the environments. *X* and *Z* are *m × p* design matrices of additive and dominance marker effects, respectively, and *β*_0_ and *θ*_0_ are their common marker effects. As reported in the SE approach (Model 1 and 2), dominance and additive versions are also tested by omitting or the *X β*_0_ or *X θ*_0_ the component in Model 4. The same standard assumptions of the GBLUP model were considered.

#### 2.3.3 Multi-environment (ME) model

The ME model is a hybrid between SE and AE approaches that includes both these models as special cases. In this application, marker effects are separated in two components: (i) a main effect estimated across all the environments, and (ii) a specific effect computed for each environment. In matrix notation, the ME additive+dominance model is expressed as following:

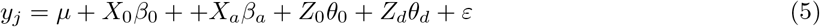

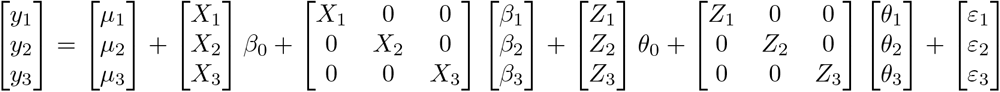

where the main additive and dominance effects are represented by *β*_0_ and *θ*_0_, respectively; specific additive and dominant effects are represented by *β_a_* ∈ { *β*_1_, *β*_2_, *β*_3_} and *θ*_a_ ∈ { *θ*_1_, *θ*_2_, *θ*_3_}, respectively. *X_a_* and *Z_a_* refer to design matrices associated to additive and dominance specific effects, respectively. Other components in the Model 5 were previously described. As reported in SE and AE approaches, ME models in their dominance and additive versions were also tested by omitting the respective terms. Likewise, standard assumptions of normality regarding the additive (common and specific effects) and dominance (common and specific effects) were considered. For the residual term, it is assumed a normal distribution with mean zero and heterogeneity of residual environmental variances, such that, *ε ~ N*(0, ∑ ⊗ *I_n_*) where ∑ is a diagonal matrix of variance-covariance structure denoting a residual variance for each environment and *I_n_* is an *n*-dimensional identity matrix.

### 2.4 Computational Implementation

The aforementioned models were implemented using the R package Bayesian Generalized Linear Regression (BGLR, Pérez and de los Campos [2013]). Common and specific marker effects were assigned as Gaussian priors (Bayesian Ridge Regression, equivalent to the GBLUP model), while flat priors were considered for the intercept. Error variances were assigned weakly informative, assuming an inverse Chi-squared density. The hyperparameters were set using the default rules implemented in the BGLR software. All models were fitted considering Markov Chain Monte Carlo (MCMC) using the Gibbs sampler with 30,000 iterations, with a burn-in of 3000, and a thinning of five. Further details related to computational implementation are described in Pérez and de los Campos [2013].

### 2.5 Assessing Model Performance

We considered two criteria to assess model performance: (i) goodness-of-fit statistics, via deviance information criterion (DIC); and (ii) predictive ability measured by cross-validation. Goodness-of-fit statistics were computed based on full data analyses. DIC is defined as a function of the deviance (likelihood function) and effective number of parameters [Gelman et al., 2014]. Models with smaller DIC values are preferred to models with large DIC. For cross-validation (CV) analyses, we considered important aspects faced by breeders when multi-environment data sets are considered (Figure 1). Two major scenarios were here defined as: i) confined prediction and ii) cross-prediction.

- **Confined prediction:** simulates a situation where GS is implemented in a specific environment, where individuals have already been genotyped and phenotyped (Fig 1a). In this scenario, marker effects were estimated using the SE model and predictive abilities were assessed using a Replicated Training-Testing evaluation [Crossa et al., 2013, Ferrão et al., 2018]. In each replication, 80% of the individuals were assigned randomly for training data set (TRN), while the remaining 20% were assigned for testing data set (TST). This division was replicated 30 times with independent random assignments into TRN and TST.
- **Cross-prediction:** simulates the question if marker effects estimated in one set of environments are useful to predict the genotype performance in another environment. In particular, we created the follow strategies: i) SE models are calibrated in a specific environment and tested in others (Fig 1a); and ii) environments were grouped to compose a new TRN data set and marker effects were estimated using AE or ME models (Fig 1a).

**Figure 1:**
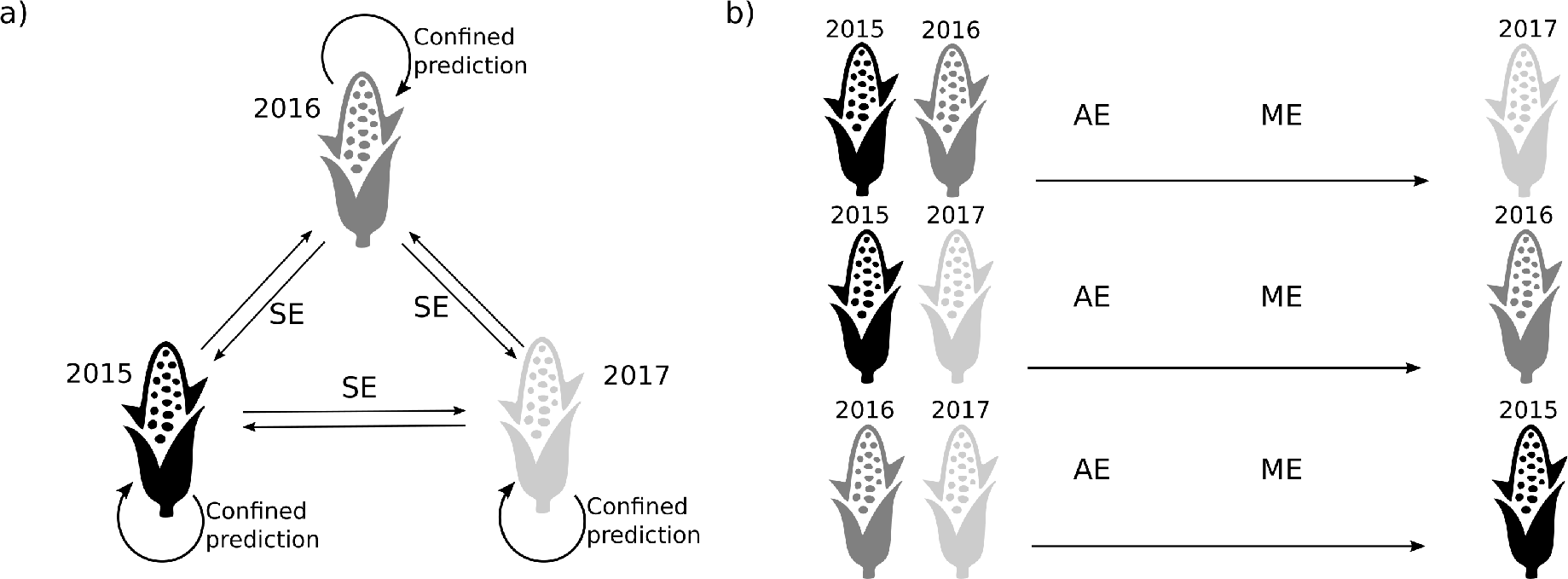
Genomic selection scenarios. a) Confined prediction represents GS implementation in a specific environment and within-sample data samples are considered in a cross-validation scheme; SE (single environment) model mimics a cross-prediction scenario when models are calibrated in a specific environment and tested in other. b) Environments were grouped to compose a new TRN data set. The estimated marker effects were used to predict a new environment using the AE (across-environments) and ME (multiple-environments) models. Direction of the arrows represented changes on training and testing data sets.

For each CV scheme, predictive abilities for the models were estimated by Pearson’s correlation between genomic estimated genetic values (GEGVs) and the corresponding phenotypic values corrected for environmental and experimental effects. GEBVs were computed as described in Table 2. For the ME models, we tested the follow alternatives to compute the GEBVs: i) considering only the common effects; or ii) considering the common effect summed to the specific effects of one of the environments.

**Table 2:** Equations used to compute the genetic merit of individuals considering the single-environment (SE), across-environment (AE) and multiple-environment (ME) models in their additive (a), dominance (d) and additive+dominance (a+d) versions. Refer to Section 2.3 for an overview of the methods compared.

| Model | Additive (a) | Dominance (d) | Additive+Dominance (a+d) |
| --- | --- | --- | --- |
| SE | $\hat{y}_{ij} = \sum_i^m X_{ij} \hat{\beta}_{ij}$ | $\hat{y}_{ij} = \sum_i^m Z_{ij} \hat{\theta}_{ij}$ | $\hat{y}_{ij} = \sum_i^m X_{ij} \hat{\beta}_{ij} + \sum_i^m Z_{ij} \hat{\theta}_{ij}$ |
| AE | $\hat{y} = X \hat{\beta}_0$ | $\hat{y} = Z \hat{\theta}_0$ | $\hat{y} = X \hat{\beta}_0 + Z \hat{\theta}_0$ |
| ME common | $\hat{y} = X_0 \hat{\beta}_0$ | $\hat{y} = Z_0 \hat{\theta}_0$ | $\hat{y} = X \hat{\beta}_0 + Z \hat{\theta}_0$ |
| ME all | $\hat{y} = X_0 \hat{\beta}_0 + X_a \hat{\beta}_a$ | $\hat{y} = Z_0 \hat{\theta}_0 + Z_d \hat{\theta}_d$ | $\hat{y} = X_0 \hat{\beta}_0 + X_a \hat{\beta}_a + Z_0 \hat{\theta}_0 + Z_d \hat{\theta}_d$ |

where: $i$ is the individual and $j$ is the environment. $\hat{y}$ is the genomic expected genetic values (GEBVs); $X$ or $X_0$ , and $X_{ij}$ or $X_a$ are design matrices for the common and specific additive effects, respectively; $Z$ or $Z_0$ , and $Z_{ij}$ or $Z_d$ are design matrices for the common and specific dominance effects, respectively; $\hat{\beta}_0$ , $\hat{\theta}_0$ , $\hat{\beta}_{ij}$ or $\hat{\beta}_a$ , $\hat{\theta}_{ij}$ or $\hat{\theta}_d$ are the common and specific marker effects estimated for the additive and dominance parametrization, respectively.

## 3 Results

### 3.1 Phenotypic Data

Figure 2 summarizes the phenotypic dispersion across the years and, for both traits, the empirical distribution was reasonably symmetric. On average, 2015 was the most productive year for grain yield and showed the largest variation. Conversely, the lowest average yield was observed in 2016. Breeding records indicated that hybrids tested in 2016 across multiple locations were submitted to abiotic stress condition caused by a reduction on fertilizing and drought, which could explain the reduced yield compared to 2015 and 2017. Grain moisture distribution is skewed to the left with heavy tails (Figure 2). On average, 2016 showed the highest values, whereas 2015 and 2017 showed similar ranges in the phenotypic distribution.

**Figure 2:**
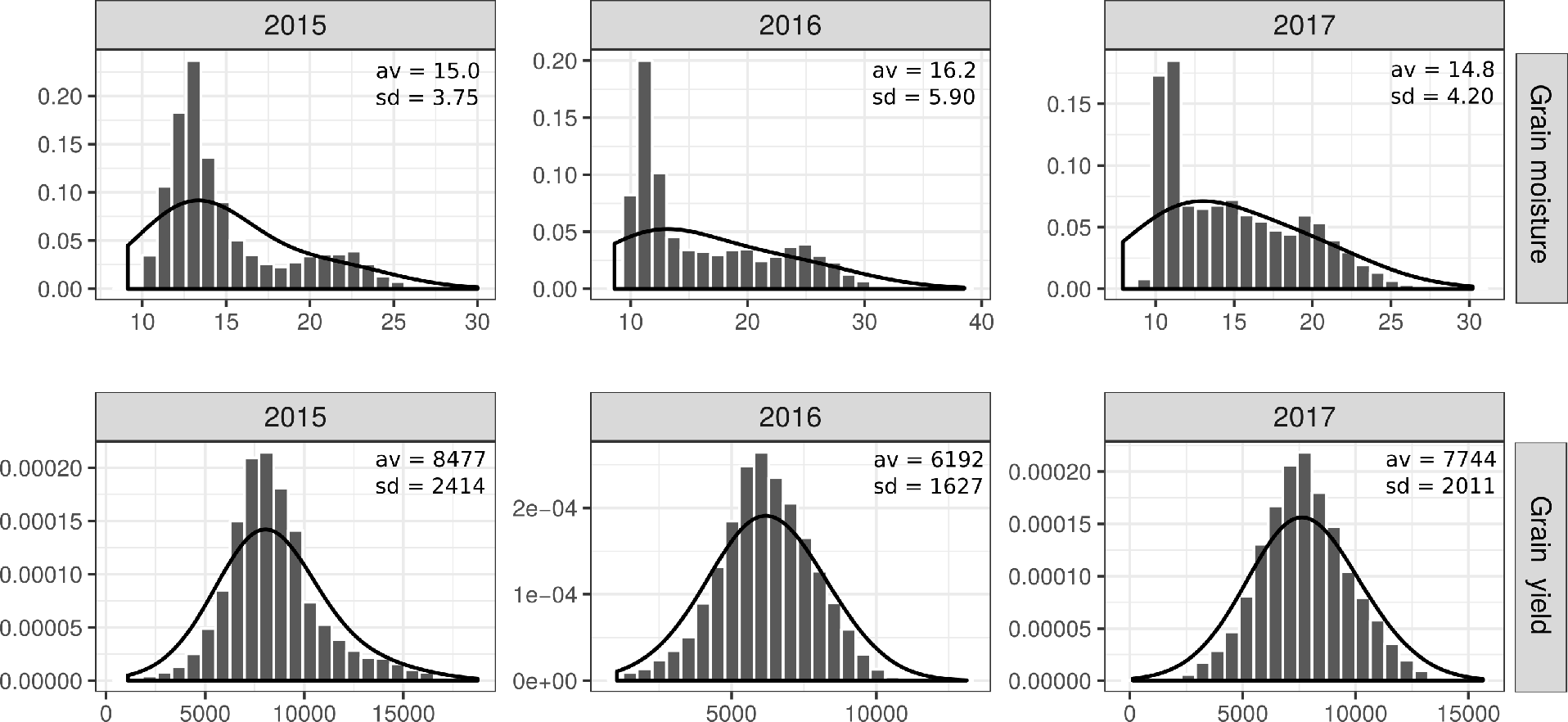
Phenotypic variation of grain moisture (%) and grain yield (Kg/ha) by environment (2015, 2016 and 2017). Average (av) and standard deviations (sd) are presented on the top of each plot.

We used multiple commercial hybrids that were planted in all locations and all three environments (years) to compute the empirical phenotypic correlation (Table 3). For both traits, correlations among environments were moderately positive. For grain production, low correlation values were observed between 2016 and other environments (Table 3, below the diagonal). A similar result was observed for grain moisture trait (Table 3, above the diagonal), which supported the observation that 2016 was an atypical evaluation cycle with a less intensive agricultural management practice.

**Table 3:** Sample correlation between grain production (below the diagonal) and grain moisture traits (above the diagonal) evaluated among the environments considering the genotype checks (commercial hybrids).

| Environment | 2015 | 2016 | 2017 |
| --- | --- | --- | --- |
| 2015 | - | 0.37 | 0.44 |
| 2016 | 0.1 | - | 0.31 |
| 2017 | 0.46 | 0.33 | - |

### 3.2 Diversity and Heterotic Groups

In this study, we evaluated a total of 1831 hybrids produced from the crosses among 207 inbred lines. Figure 3a represents the maize hybrids that were tested in the field and illustrate the practical challenge in testing all inbred combinations, given the demand of large experimental area and labor. The importance of GS lies in the possibility to accurately predict hybrids that were not phenotyped - represented in Figure 3a by white spots. The genomic data indicated the presence of 3 clusters/subpopulations that may represent the number of heterotic groups in the current germplasm collection (Figure 3b and 3c). PCA and k-means analysis estimated three main groups and low overlap between them. Definition of the number of clusters was based on the dramatically reduction observed in the SSE values.

**Figure 3:**
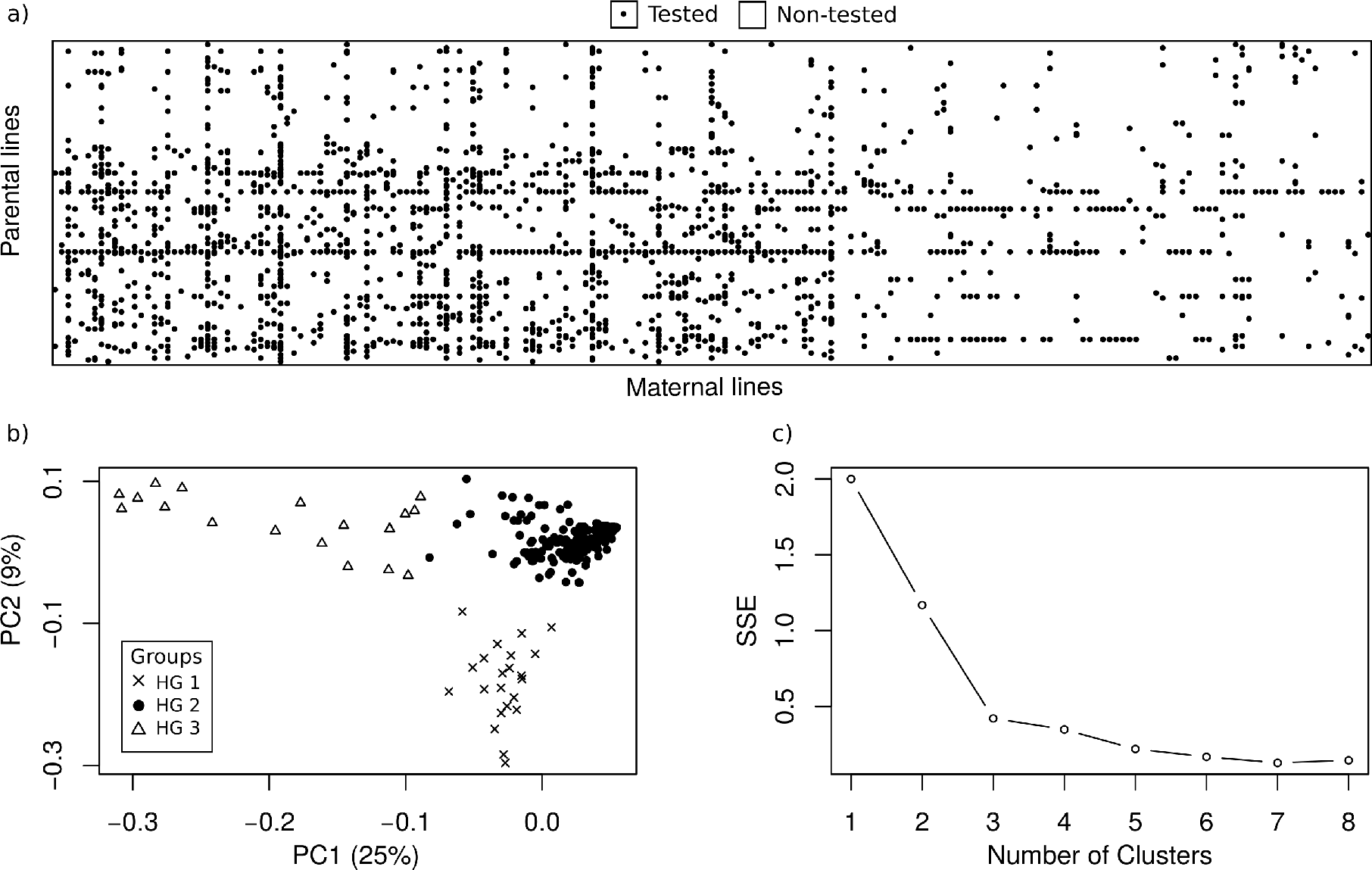
a) Schematic representation of tested hybrids obtained by crossing parental with maternal lines. Hybrids tested in the field are represented by black circles, while the white spots are representing hybrids not phenotyped. b) Plot of the two eigenvectors of the genomic relationship matrix for the lines. c) Sum of squared error (SSE) for a number of cluster solutions.

### 3.3 Goodness-of-fit statistic, Variance Components and Predictive Ability

For both traits, the lowest DIC values were obtained for additive+dominance and multi-environment models, which evidence the importance to simultaneously account for dominance gene action and G*×*E interaction (Table 4). A summary of predictive performance for grain yield and grain moisture are presented in Figure 4, 4 and 6. Results can be described under two different perspectives: i) importance to include dominance effects in GS models; and ii) assessment of predictive performance considering different CV schemes.

**Table 4:** Goodness-of-fit value for grain yield and grain moisture traits for across-environments (AE) and multiple-environments (ME) models in their additive, dominance and Additive + Dominance versions.

| Model | Grain Yield |  | Grain Moisture |  |
| --- | --- | --- | --- | --- |
|  | AE | ME | AE | ME |
| Additive | 6233.6 | 6032.3 | 4565.4 | 4315.6 |
| Dominance | 5735.8 | 5329.3 | 4641.6 | 4256.5 |
| Additive + Dominance | 5731.7 | 5292.9 | 4512.8 | 4083.5 |

In view of gene action, the results are better illustrated in Figure 4. Predictions performed within-environment and accounting for dominance effects were more accurate for grain yield. Conversely, additive models were slightly better than dominance models and equivalent to additive+dominance for grain moisture (Fig 4a). Prediction accuracies ranged from 0.34 to 0.67 for grain yield and 0.67 to 0.82 for grain moisture. The difference in performance between additive and dominance models was smaller for grain moisture than for grain yield (average difference of 0.02 vs. 0.16). In almost all scenarios, modelling simultaneously both gene actions did not impact the predictive performance. Based on the goodness-of-fit values, we considered the most parsimonious models to quantify the role of each genetic component on the phenotypic variation (Fig 4b). The residual variance estimates varied across environments (Fig 4b) suggesting that homogeneous residual variance assumed in the AE model may not be optimum in this context. Finally, the proportion of variance explained for each term was estimated for the ME additive+dominance GBLUP. Observed results were in line with the predictive abilities, where the variance explained by the dominance effects in grain yield was greater than additive effects; whereas for grain moisture the result was the opposite

**Figure 4:**
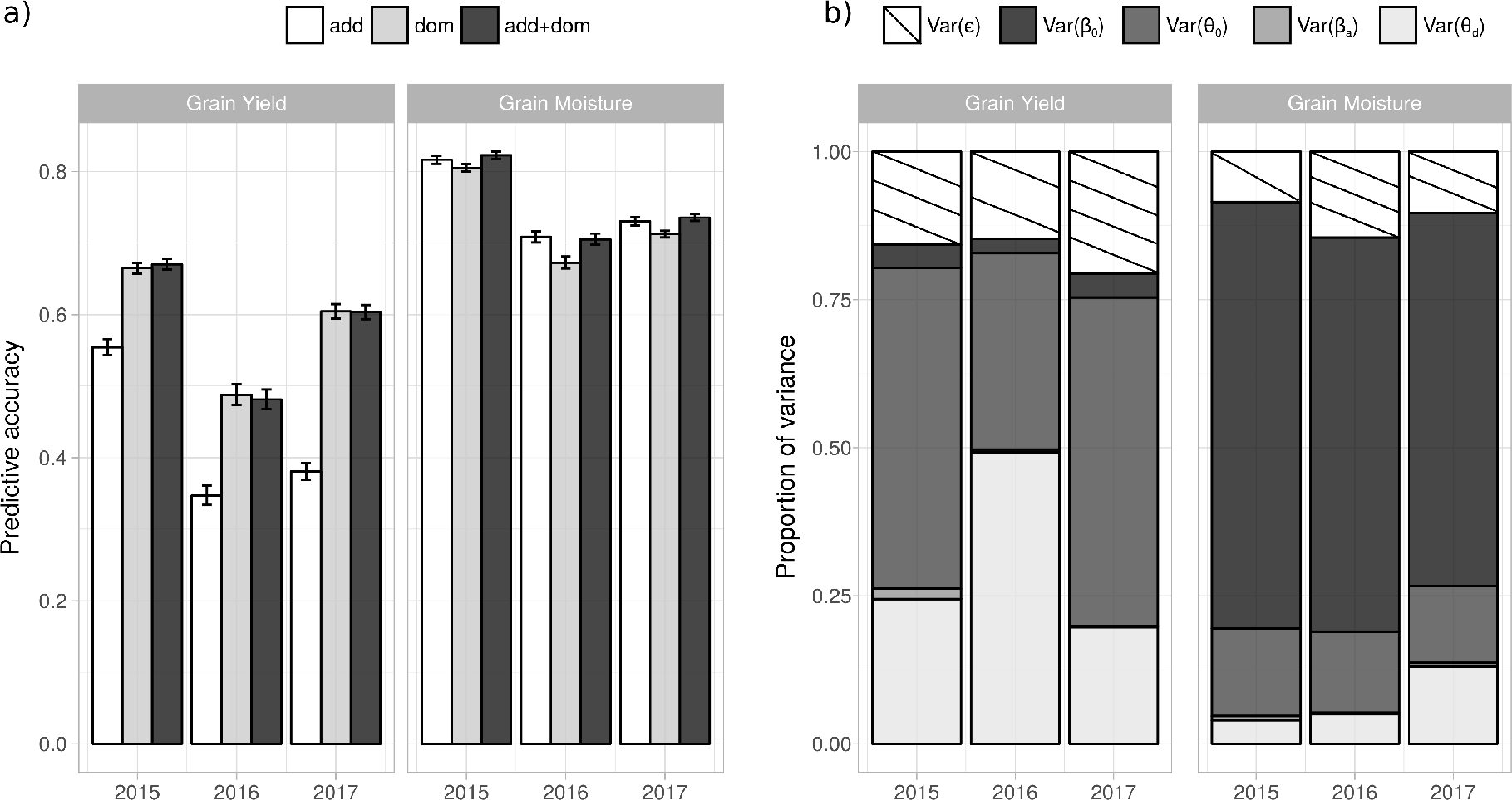
a) Predictive accuracy of within-environment models using single-environment (SE) models for grain yield and grain moisture traits evaluated in 2015, 2016 and 2017. All models were considered in their additive (add), dominance (dom) and additive+dominance (add+dom) versions. Refer to Section 2.3 for an overview of the methods compared. b) Estimates of variance components considering the additive+dominant version of the multiple-environment (ME) model grain yield and grain moisture traits evaluated in different 2015, 2016 and 2017.

In terms of breeding schemes, for grain yield, the highest performances were observed in confined predictions (Fig 5a). Under this circumstance, models trained and validated in 2015 showed the highest predictive performance (0.67, for the additive+dominance version). Confined prediction performed in 2016 showed the lowest values (0.35, for the additive model), while confined predictions in 2017 yielded intermediate values, closer to those observed in 2015 (0.60, for the additive+dominance version). Despite differences in population size, this feature was not the primary reason to affect the prediction accuracy (e.g., 2016 yielded lower performance than 2015, even with a larger population size). Predictions performed in new environments were investigated considering cross-prediction schemes. In this scenario, we observed variable performance over the strategies tested. Single environment analysis (SE models) proved to be highly dependent on the TRN-TST setting. For example, assuming the additive+dominance model for predictions in 2017, predictive ability ranged from 0.56 to 0.16 when the models were trained on either 2015 or 2016, respectively (Fig 5a). In contrast, considering the AE and ME models to predict 2017, the values ranged from 0.29 to 0.57 (Fig 5b). Compared to the worst scenario observed in the stratified analysis, this value represents an increase of 55% in predictive performance.

**Figure 5:**
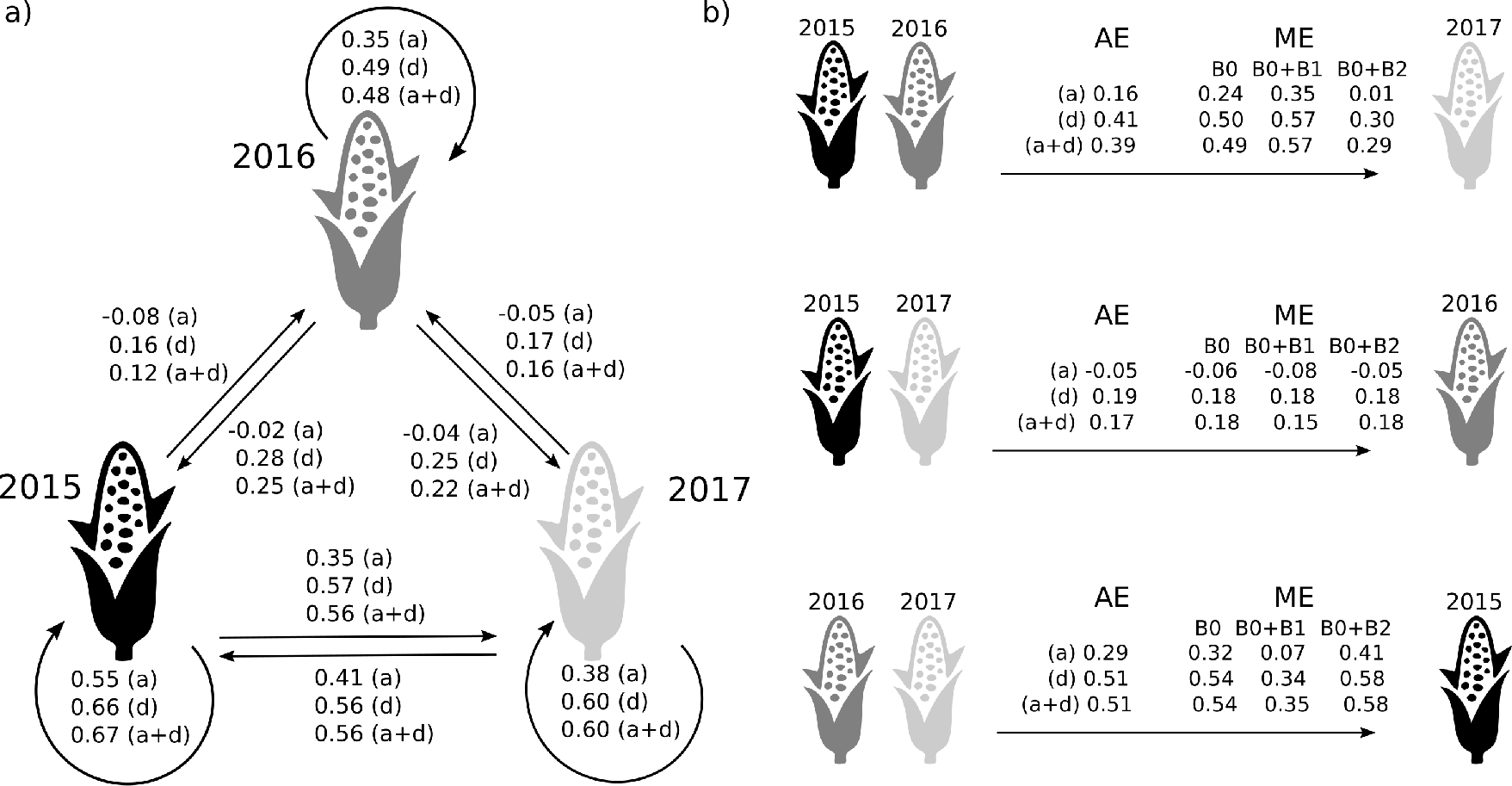
Correlation between phenotypes and predictions for the grain yield trait in maize. a) Predictive performance assessed within the same validation scheme (average over 30 TRN-TST partitions) and considering stratified analysis (models training in an environment and tested in other). b) Predictive performance considering combined environments assuming the across-environment (AE) and multi-environment (ME) modelling. For the ME modelling, predictions were performed considering only the main effect (B0, equivalent to *β̃*_0_ and *θ̂*_0_) and the sum of the main with a specific effect (B1 and B2, *β̂*_*a*_ and *θ̂*_*d*_, that represent the environments considered in the TRN partition in the same order presented in the figure). Refer to Section 2.3 for an overview of the methods compared.

In contrast, for grain moisture, on average, higher values of predictive performance were observed. Similarly, the highest predictive value was observed in confined predictions (Fig 6); where the environment 2015 showed the best performance (0.82, for the additive+dominant model), followed by 2017 (0.73, for the additive+dominance model) and 2016 (0.70, for the additive+dominance model). Cross predictions over the environments also showed variable performance. Unlike grain production, models trained in 2016 did not result in low predictive performance when validated in 2015 and 2017. Accordingly, the use of combined environments and ME models showed more consistent results irrespective of the environments used in the TST set.

**Figure 6:**
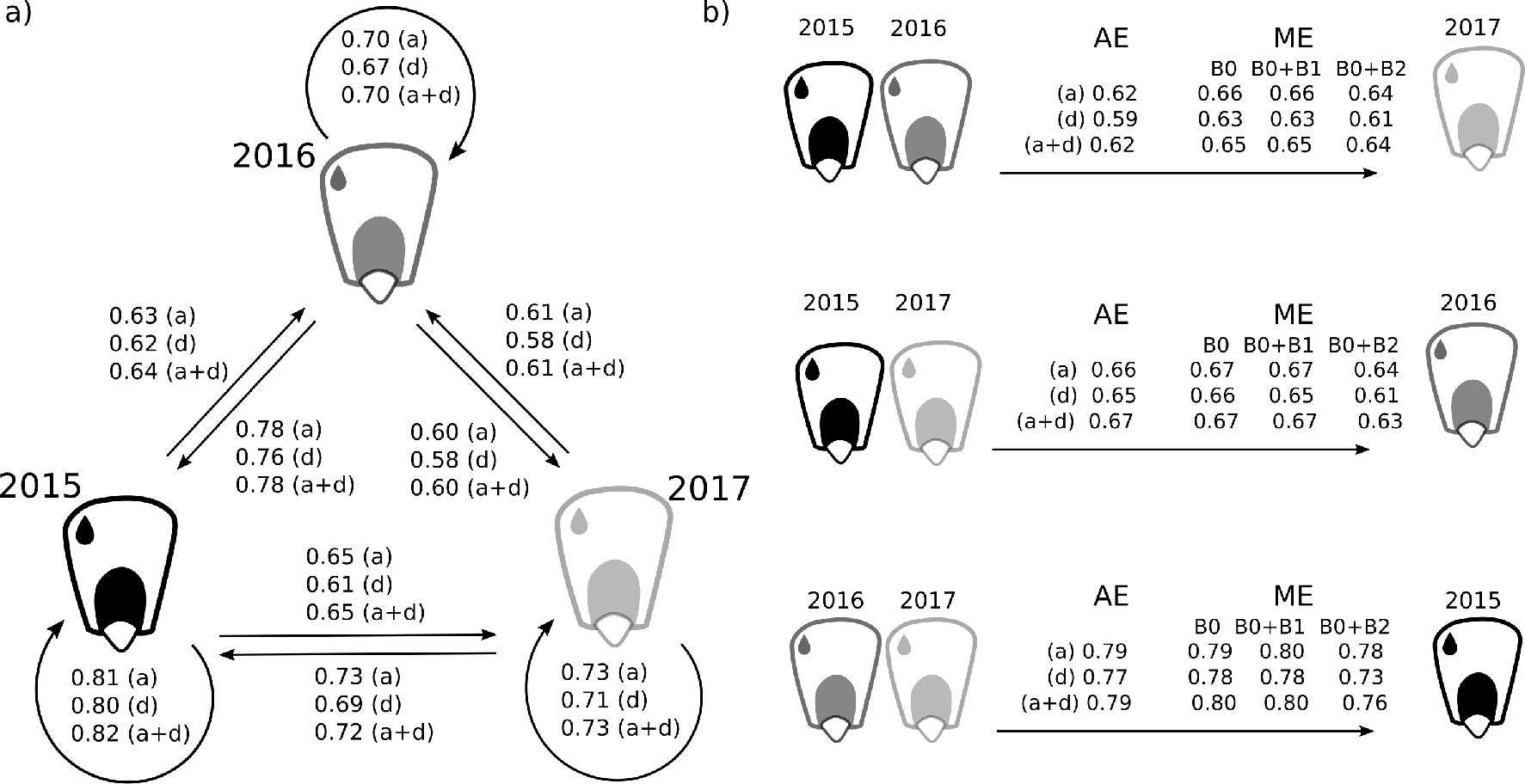
Correlation between phenotypes and predictions for the grain moisture trait in maize. a) Predictive performance assessed within the same validation scheme (average over 30 TRN-TST partitions) and considering stratified analysis (models training in an environment and tested in other). b) Predictive performance considering combined environments assuming the across-environment (AE) and multi-environment (ME) modelling. For the ME modelling, predictions were performed considering only the main effect (B0, equivalent to *β̂*_0_ and *θ̂*_0_) and the sum of the main with a specific effect (B1 and B2, *β̂*_*a*_ and *θ̂*_*d*_, that represent the environments considered in the TRN partition in the same order presented in the figure). All models were considered in their additive (a), dominance (d) and additive+dominance (a+d) versions. Refer to Section 2.3 for an overview of the methods compared.

As previously cited a potential benefit to explicit modelling M*×*E interactions is to decompose marker effects into components that are stable and specific across environments. Naturally, this approach could be used to identify genomic regions that have constant effects across groups and ones that exhibit substantial interaction on the phenotypic variation. To illustrate this, Figure 7a shows how the common marker effects (*β̂*_0_ and *θ̂*_0_) estimated under the additive and dominance ME models are ranging along the genome. Differences in the magnitude values are evidencing the role of gene action conditional to the traits. In Figure 7b, we illustrate the range of additive (*β̂*_1_ and *β̂*_2_) and dominance (*θ̂*_1_ and *θ̃*_2_) marker effects estimated in 2015 and 2016 for grain yield and grain moisture, respectively. The results suggest differential effects across the years.

**Figure 7:**
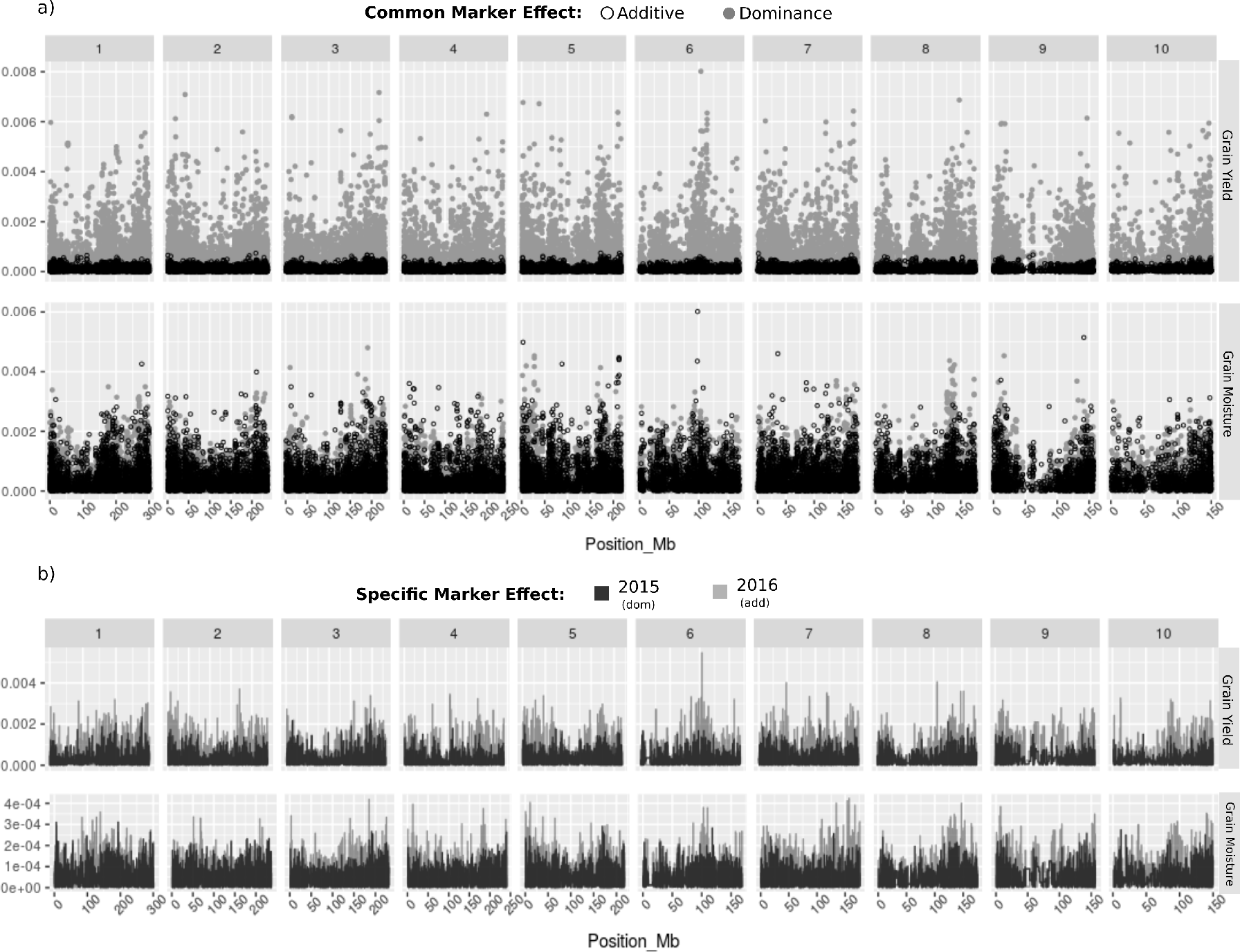
Correlation between phenotypes and predictions for the grain moisture traia) Manhattan plot for grain yield trait and grain moisture evaluated in maize hybrids. Black and gray dots are representing, respectively, additive (*β̂*_0_) and dominance (*θ̂*_0_) common effects estimated using the multiple-environment (ME) model across three years of evaluation (2015, 2016 and 2017). b) Profile of dominance (*θ̂*_1_ and *θ̂*_2_)and additive (*β̂*_1_ and *β̂*_2_) marker effects estimated for grain yield and grain moisture traits considering the specific marker effects estimated under the ME model in 2015 and 2016. Refer to Section 2.3 for an overview of the methods compared.

## 4 Discussion

Progress in hybrid breeding can be greatly accelerated by the incorporation of genomic predictions into breeding schemes [Technow et al., 2014, Acosta-Pech et al., 2017, Dias et al., 2018, Werner et al., 2018]. To this end, breeders have to face important issues regarding its implementation, including the impact of accounting for non-additive effects and dealing with G*×*E interaction. In this work, we report the effects of these issues in the predictive performance, definition of training population and allocation of resources in a breeding program.

From classical quantitative genetics theory, dominance effects are defined as intra-locus interactions resulting from differences between the genotypic value and the breeding value [Falconer and Mackay, 1996, Lynch and Walsh, 1998]. In GS modelling, a common practice has been to ignore it under the argument that the additive component is larger and therefore more important than dominance effects [Huang and Mackay, 2016]. Although some results have been supporting this evidence [Hill, 2010], renouncing the importance of dominance effects may be at least controversial if we consider that “dominance” and “overdominance” are two of the most accepted theories on the genetic basis of heterosis.

In this study we have demonstrated that the inclusion of dominance effects increases the predictive ability, in particular, for grain yield. Notoriously, in some cross-validation scenarios the predictive capacity increased by 30% when compared to additive models. Our findings are also supported by goodness-of-fit values since dominance models are more parsimonious than the additive model. Accordingly, other studies have shown similar empirical results [Resende et al., 2017, Dias et al., 2018]. Despite the importance of dominance effect in grain yield, additivity explained a large portion of variance in grain moisture, suggesting that both traits have different genetic architectures. Nonetheless, the inclusion of dominance into additive+dominance models resulted in equivalent and sometimes better predictive abilities compared to additive models. Therefore, one important contribution of this study is to demonstrate that, regardless of the underlying genetic architecture of a trait, considering both gene action in GS models is a valid alternative to achieve high prediction performance.

Considering the gene action modelling, another related issue is the difficulty to properly separate additive and dominance effects in genetic analysis [Huang and Mackay, 2016]. At least as conventionally applied, orthogonal partitions are achieved only assuming theoretical conditions. For example, in the classical model proposed by Fisher [1918] and developed by Cockerham [1954] and by Kempthorne [1954], orthogonality assumptions were derived assuming Hardy-Weinberg equilibrium. Although it provides an elegant formalization of the genetic variance partition, it is well-known that many precepts assumed are not applied to artificial populations [Hill, 2010]. Herein, we have reported a simple parametrization of dominance, which may be biased by non-orthogonality of genetic effects. Despite the apparent severity to induce erroneous interpretations, our conclusions on the importance to account for dominance effect are primarily supported by the improvement of the predictive results.

A second contribution of this investigation is to discuss the GS implementation, with a particular focus on G*×*E modeling. Many studies have been shown the high performance of GS in single environments. Herein, we defined this strategy as confined predictions and, for both traits, high values of predictive ability were observed. Accordingly, Technow et al. [2014], Acosta-Pech et al. [2017] and Zhao et al. [2012] reported similar results in maize. Biologically, it is reasonable to expect higher predictive values when CV schemes are setting using within-sample data, since training and validation data set are exposed to the same environmental inputs. Despite the high performance, this scenario does not attempt to answer an important question in a breeding program that involve predictions in a new set of environments. If feasible, marker effects estimated in one set of environments would be considered to predict the genotype performance in a different environment, without the necessity of retraining the models. This immediately suggests a reduction of time and resources spent in field evaluations [Resende et al., 2012, Ferrão et al., 2018].

To assess predictive performance of new environments, we tested three approaches: single-environment (SE), across-environment (AE) and multiple-environment analyses (ME). SE and AE models are based in opposed assumptions. Precisely, SE modelling assumes a regression model for each single environment, where none of the information from different environments is combined. In contrast, AE model considers a common marker effect and hence a regression model is fitted for the combined data set, unaware that the data came from different environments. The relative performance of these models would therefore be expected to vary depending on the G*×*E magnitude. This is evident when SE model is trained in 2015 and validated in 2017, for grain production. Based on the phenotypic correlations, both environments are considered similar, which suggests less noise caused by the G*×*E interaction. As result, high predictive ability was observed under the SE model. However, this same model is not the most efficient to predict 2016. As evidenced by the phenotypic correlations, 2016 is an atypical environmental condition and thus a contrasting environment. Consequently, SE models did not yield high predictive values and, predictions performed in 2016 are benefited by the combination of 2015 and 2017 considering the AE model.

Differences in predictive performance across the methods are demonstrating that focusing on one approach may adversely affect the final result. In practice, one does not know the properties of a new environment and how related that new environment is any testing set. Thus, it is unclear which of the two models (SE or AE) should be considered. Assuming that ME is a hybrid between SE and AE models, it naturally has a wide range of potential uses. As a consequence, we noted that ME is either as accurate, or more accurate, than the two competing models presented in this work. Predictions performed in 2017 for grain yield provides a good example. The highest values in this year are achieved using the ME model, when specific effects estimated in 2015 are accounted with the common effect. On the other hand, poor results are observed when specific effects from 2016 are considered. In situations, which many environments are considered and the G*×*E is unknown, defining which specific effect should be weighted with the common effect may be complicated. A conservative alternative would be to consider only the common effect estimated by the ME model. Similarly, results reported in wheat and maize are supporting the importance of ME models [Lopez-Cruz et al., 2015, Crossa et al., 2016, Cuevas et al., 2016a,b, Crossa et al., 2017].

Despite our focus in genomic prediction results, the use of molecular markers in maize breeding has promising applications. We briefly consider its potential to define heterotic group in our current germplasm. Historically, our heterotic group classification has been based on pedigree records and visual characterization, which commonly result in errors. According to breeders, the groups suggested in this research are in accordance to historical records, which point out the importance to use molecular markers to access the genetic diversity. Considering the ME modelling, our results suggest its potential to genome-wide association studies (GWAS), since marker effects on grain yield were individually estimated. We emphasize that identified regions harboring SNPs that affect some phenotype or outcome of interest is a goal that can naturally be cast using this approach. More recently, many GWAS investigations described similar procedures that fit all SNPs simultaneously as random effects, which approximate GWAS and GS models [Guan and Stephens, 2011, Karkkainen and Sillanpaa, 2012, Goddard et al., 2016].

Finally, we hope that our work also helps highlight some general guidance for practical GS implementation. Based on our results we proposed: i) compute GEBVs considering additive and dominance effects for different traits; ii) use of multi-environment models to predict genotypic performance in a new environment; iii) refine our heterotic group classification based on molecular information; and iv) use SNP data to, *in silico*, extrapolate maize hybrid composition and, based on genome predictions, select the best genotypes for breeding. All proposed approaches are computationally tractable for moderately large datasets and are flexible to perform well in a wide range of conditions. Furthermore, results interpretation and computational implementation are straightforward. We also emphasize that our study is a first stage of what could be done considering dominance and multiple environment models in GS studies. In particular, we believe that our results are sufficiently promising to justify further research, including the test of different parameterizations for dominance effects and incorporation of multiple environmental covariates for G*×*E modelling.

## 6 Author Contributions

MFRRJ coordinated the study. CDM conducted the field experiment and collected the genotypic and phenotypic data. LFVF, CDM and MFRRJ performed the data analyses and interpretation. LFVF and MFRRJ wrote the paper. PRMV provided expertise and edited the manuscript. All authors read and approved the final version of the manuscript for publication.

## 7 Acknowledgments

We sincerely thank Dr. Gustavo de Los Campos for providing useful comments and suggestions on this manuscript. We also thank the company Helix Seeds to provide the data set used in this investigation.

